# Low Effective Dimensionality Does not Invalidate Symptom Lesion Network Mapping

**DOI:** 10.64898/2026.07.29.741272

**Authors:** Thomas Zaugg, Vladimir Litvak

## Abstract

Van den Heuvel et al. argued that the low effective dimensionality of the human connectome causes lesion network maps to converge on a common pattern, the connectome degree map, and concluded that the method cannot resolve distinct symptom-specific circuits. Here, we show that, although lesion network maps are low-dimensional and biased towards the connectome’s leading eigenvector, they still recover their ground-truth networks well. Moreover, we propose a modification of the connectome that at least halves this bias and can improve recovery. Our results, therefore, reaffirm the validity of lesion network mapping, while acknowledging the important limitations identified by its critics.

---

Lesion network mapping (LNM) has gained prominence as a method for relating sets of non⍰overlapping brain lesions to the symptoms they produce. LNM maps symptom-specific functional circuits and suggests potential targets for invasive and non-invasive neuromodulation.

Van den Heuvel et al. recently argued that the low effective dimensionality of the human connectome leads lesion network maps derived from any set of lesions to converge on a common pattern, namely the connectome degree map^1^. They therefore concluded that LNM cannot resolve distinct symptom-specific circuits.

Subsequent discussion has identified limitations in this analysis and reasserted the empirical validity of LNM^2–5^. However, no formal framework has shown how LNM can remain informative despite van den Heuvel’s critique, which rests on the dominance of a small number of principal components, in particular the connectome’s leading eigenvector (PC1).

Treeratana et al. (2026) proposed a simulation to test the ideas put forward by van den Heuvel and colleagues^6^. They introduced a generative model that links lesions to symptom scores, conditional on a specific ground truth (GT) network assumed to underlie those symptoms. The set of 300 GT networks was derived from the functional connectivity patterns of parcels in the Schaefer brain atlas^7^ (see Supplementary Note 1).

The authors then evaluated the extent to which GT networks can be recovered using a symptom Lesion Network Mapping (sLNM) approach. They show that although sLNM maps recover their GT network, they simultaneously overrepresent PC1 features relative to GT, which can lead to spurious similarities between disorders. To address this, they propose to regress PC1 out before comparing maps.

Here, we extend the simulation framework of Treeratana et al. to address two questions. First, to what extent symptom-specific circuits remain recoverable despite the low-dimensional structure of the connectome and the bias towards PC1. Second, where this bias originates and how it can be corrected.

We show that, although each lesion network map is low-dimensional, it recovers its GT network well above the level achievable from PC1 alone. The distinct network features are therefore preserved under the low dimensionality of the connectome. We also confirm that sLNM maps show a systematic bias towards PC1, which inflates inter-map similarity and increases the number of significant parcels under permutation testing. We trace this bias to the PC1-dominated connectivity structure of lesion-derived maps and propose a correction that at least halves the PC1 bias and can improve recovery in the resulting sLNM maps, depending on how much PC1 bias the observed symptom scores carry.

Our conclusion, therefore, contrasts with that of van den Heuvel et al. We reaffirm the validity of lesion network mapping, while acknowledging the important limitations identified by its critics.

## Similarity structure of functional networks is preserved in low dimensions

We used three measures to quantify the similarity between two network maps X and Y:

- **spatial r**: Pearson r between X and Y (computed across parcels).
- **symptom r**: Pearson r between the symptom scores predicted by X and those predicted by Y.
- **top-100 overlap:** fraction of 100 highest-value parcels (=top-10%) of map X among the 100 highest value parcels of map Y.

Examining the similarity structure of network pairs ordered by their correlation with PC1 shows that, although the GT set was optimised to maximise distinctiveness, most network pairs remain substantially (anti-)correlated (Fig. 1a). The sLNM procedure further enhances pairwise similarity, analogous to increasing image contrast, such that the similarity between pairs of sLNM maps generally exceeds that of their corresponding GT networks. The degree of similarity is particularly high for symptom r, with most GT pairs showing strong cross-predictive performance. This is important because cross-prediction has previously been interpreted as evidence that networks are functionally identical, a concern also raised using clinical LNM maps^6^. In contrast, the top-100 overlap metric retains a more granular structure, consistent with prior suggestions that networks can remain distinct in their peak regions despite high overall correlation (Fig. 1a)^8,9^.

**Fig. 1:**
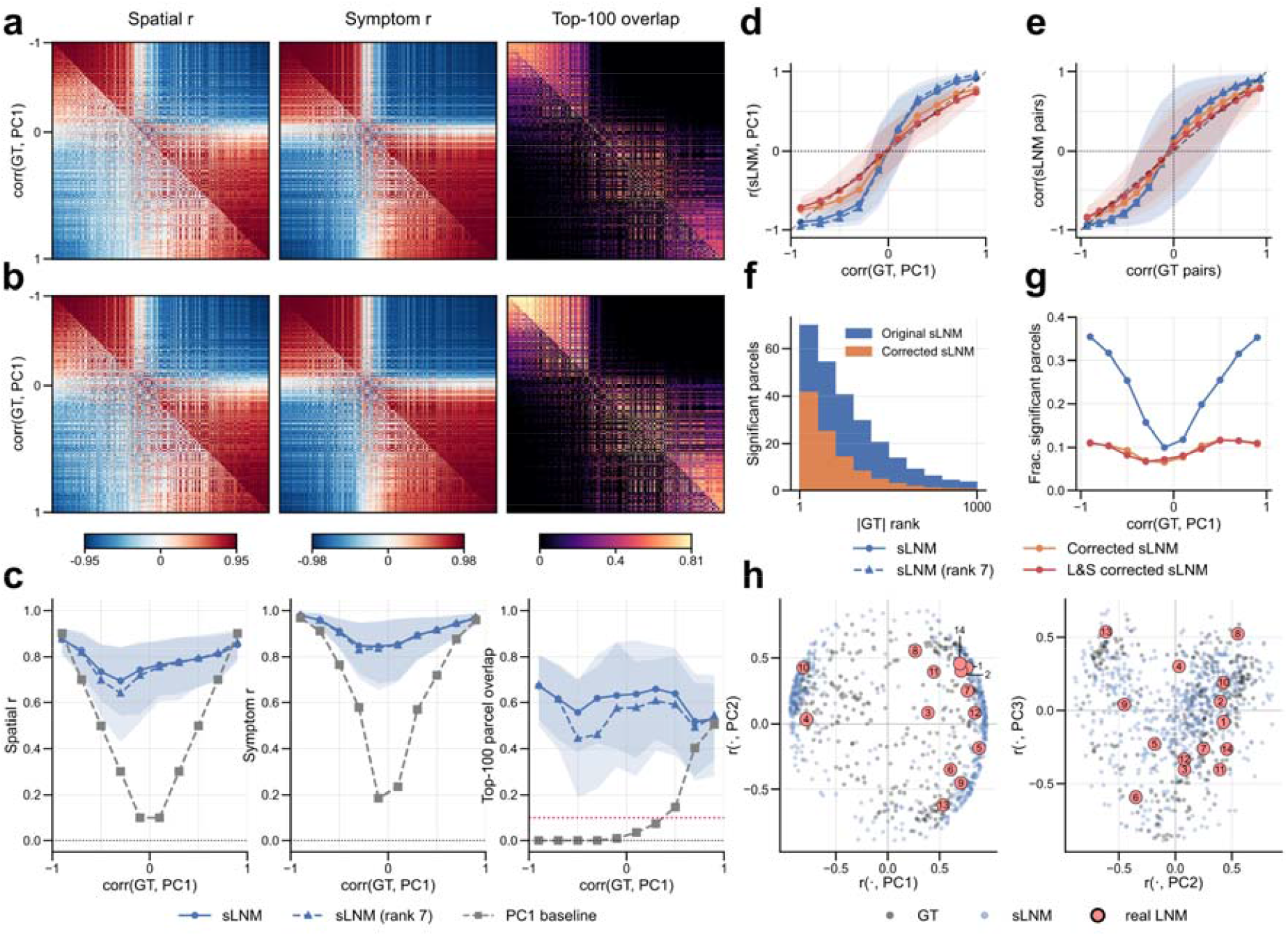
Throughout, sLNM denotes symptom lesion network maps; PC1 is the leading eigenvector of the group connectome. Conditions: *original sLNM* (blue), *rank-7 reconstruction of the original sLNM* (blue, dashed), *corrected sLNM* (orange), and Lesion & Symptom (*L&S) corrected sLNM* (red; used in d,e,g). Confidence bands span the 5th–95th percentile across 300 GT networks and 10 noise realizations each; η^2^ = 0.3 throughout. **a**,**b)** Pairwise similarity among the 300 GT networks (lower triangle) and the corresponding sLNM (upper triangle), sorted by corr(GT, PC1). *Spatial r* (left): the block structure of the GT set (lower triangle) is reproduced but inflated in the sLNM (upper triangle). *Symptom r* (middle) between map pairs is strongly positive or negative, tracking the sign of each pair’s PC1 alignment. *Top-100 overlap* (right) shows fine-grained spatial structure for both GT and sLNM. The overall pattern from the full-rank sLNM (a) is preserved in the rank-7 reconstruction (b). **c)** Recovery of GT for the original sLNM and a rank-7 reconstruction. Spatial (left) and symptom r (middle) show almost no difference for the rank-7 reconstruction, top-100 overlap (right) decreases slightly. Crucially, recovery stays above the PC1 baseline for all conditions. sLNM, therefore, carry GT-specific structure beyond their PC1 alignment. **d)** PC1 alignment of the sLNM as a function of PC1 alignment of their GT. *Original sLNM* (blue) show a high PC1 bias (deviation from the identity line), especially for moderately PC1-aligned GTs. *Corrected sLNM* (orange) show half the bias, while Lesion & Symptom (L&S) corrected sLNM (red) have essentially no bias. **e)** Similarity of sLNM pairs against similarity of their GT pairs reveals over-similarity among original sLNM (blue) pairs relative to their GT pairs, driven by PC1 alignment. Corrected sLNM (orange) show less over-similarity; L&S corrected sLNM (red) essentially none. **f)** Distribution of significant parcels by their unsigned rank in the GT, for original (blue) and corrected (orange) sLNM. Significant parcels concentrate at high ranks of the GT in both conditions, indicating that significance tracks genuine GT structure. **g)** Fraction of significant parcels as a function of corr(GT, PC1). For the original sLNM (blue) the fraction rises steeply toward both extremes; for the corrected sLNM (orange) and L&S corrected sLNM (red) the curve is flatter. The curves nearly coincide at low |corr(GT, PC1)| and diverge only toward the extremes, showing the correction removes PC1-dependent inflation in significant parcels while leaving baseline parcel detection intact. **h)** Simulated sLNM (blue), parcel-GT networks (gray) and real LNM maps (salmon) plotted in the first three principal components (PC1–PC3). *Left:* PC1 vs. PC2. *Right:* PC2 vs. PC3. GT networks are scattered throughout; simulated sLNM are more concentrated towards the PC1 poles. The 14 real LNMs remain distinguishable from one another in the first three PCs, confirming that sLNM maps retain lesion-specific information rather than collapsing onto the leading eigenvector, despite the connectome’s low effective dimensionality. Real LNM are identified in Supplementary Table 1, with corresponding references.

To test whether these patterns depend on high-dimensional details, we reconstructed each sLNM map from its first seven connectome-eigenmode coefficients (a rank-7 reconstruction), which recovers most of the variance (mean r^2^ = 0.87). Reconstructed networks retain nearly identical similarity structure on all three measures (Fig. 1b).

Despite this low dimensionality, each map recovers its own GT across all three metrics, with spatial r = 0.79, symptom r = 0.91, and top-100 overlap = 0.60. The rank-7 reconstruction achieves comparable recovery to the full map, with spatial and symptom r largely unchanged at 0.78 and 0.91, and a reduction in overlap from 0.60 to 0.55 (Fig. 1c).

This drop in overlap reflects a property of the GT network itself: building a lower-rank reconstruction of the GT shows that its top-100 parcels are only recovered using many more components than what is sufficient for spatial and symptom r (Supplementary Fig. 1). This suggests that accurate recovery of peak regions in functional maps requires higher-resolution representations of the connectome. Consequently, the simplification from voxel-based to parcel-based maps, as used by van den Heuvel et al., may lead to missed peaks.

## PC1 bias

Lesion network maps show a systematic bias towards PC1 compared with their corresponding GTs, as established by Treeratana et al.^6^. The magnitude of this bias depends non-linearly on the similarity between the GT and PC1, with moderately correlated GTs having the highest bias (see Fig. 1d). Because of this PC1 bias, any two sLNM maps are more (anti-)similar to each other than their respective GT networks are (Fig. 1e), driving the contrast-like enhancement of pairwise similarity described above (Fig. 1a).

We trace the source of this PC1 bias to the lesions’ functional connectivity (FC) maps. A lesion’s FC map is generated by passing its seed through the connectome. Since the connectome’s variance is concentrated in a few leading eigenmodes, this operation systematically amplifies those top modes relative to the rest: even a seed only modestly aligned with PC1 yields an FC map dominated by it (see Supplementary Note 2 for derivation). Since every lesion FC map is skewed toward PC1 in this way, the sLNM maps built from them inherit the same alignment.

Van den Heuvel et al. argued that over-similarity with the connectome degree map could mean sLNM maps fail to recover their specific GT^1^. However, we find that across the full range of corr(GT, PC1), sLNM recovery stays well above what PC1 alone would achieve as a predictor (Fig. 1c). sLNM maps created from GT networks that are low or only moderately correlated with PC1 still recover their GT, which a purely PC1-driven map could not. sLNM maps, therefore, carry GT-specific structure beyond PC1.

## Corrected sLNM

We propose a simple correction for this PC1 bias: constructing the lesion FC map from a connectome whose leading eigenvalues have been flattened. We refer to maps built from these corrected lesion FC inputs as *corrected sLNM maps* (see Supplementary Note 3).

*Corrected sLNM maps* show roughly half the PC1 bias of the original maps (mean |r(sLNM,PC1) − r(GT,PC1)| 0.23 → 0.12; Fig. 1d), with a corresponding reduction in inter-map over-similarity (mean |r(sLNML□,sLNML□) − r(GTL□,GTL□)| 0.36 → 0.23; Fig. 1e). The residual bias enters through the symptom scores, which in the simulation are computed from the original PC1-dominated lesion FC maps. Recomputing the symptom scores from corrected FC maps (*Lesion & Symptom corrected sLNM*) further removes most of the remaining PC1 bias (Fig. 1d), leading to essentially no over-similarity (Fig. 1e). This isolates the two routes by which the PC1 bias enters the sLNM maps in our simulation framework: the FC maps used to build the sLNM maps, and the FC maps used to generate the symptoms.

In real applications, symptom scores are observed clinical data, not computed from lesion FC maps, so only the lesion-side correction is deployable. After eigencorrection of the lesions, any residual PC1 bias in sLNM maps is therefore inherited from the symptom scores. Whether this residual symptom bias exists, and whether it reflects genuine structure in the symptom-connectome relationship, as suggested by Zalesky and Cash^8^, cannot be determined from lesion data alone and remains an open empirical question.

The eigencorrection of the lesions alone (*corrected sLNM maps*) improves recovery for GTs weakly or moderately correlated with PC1, while recovery for strongly PC1-aligned GTs falls slightly (Supplementary Fig. 2c); the latter is expected, since the original method’s PC1 bias inflated recovery because PC1 approximates the GT in these cases. The net effect is a recovery profile that is more constant across *corr(GT, PC1)*, with aggregate recovery essentially unchanged (spatial r: 0.79 → 0.78; symptom r: 0.91 → 0.93; top-100 overlap: 0.60 → 0.58).

The *Lesions & Symptoms (L&S) corrected sLNM maps* give an upper bound on what the lesion-side correction could achieve if empirical symptoms carry little residual PC1 bias of their own (spatial r: 0.79 → 0.82; symptom r: 0.91 → 0.94; top-100 overlap: 0.60 → 0.66; Supplementary Fig. 2c). The true benefit of the lesion-side correction on real data therefore lies somewhere between these two correction profiles, depending on how much PC1 bias the symptoms carry. In either case, the excess PC1 bias in the original sLNM map is an artifact of the PC1-dominated lesion FC map that can be substantially reduced.

## Permutation testing

Permutation testing is used to control for spurious LNM findings. We used a symptom-permutation null with family-wise error control^4,8^. For each permutation, we record the maximum absolute parcel value across all 1000 parcels; the 95th percentile of this max-statistic distribution sets a single threshold that controls FWER across the whole map (see Supplementary Note 4 for differences with Treeratana’s null).

Under the η^2^=0 (no-signal) control, every significant parcel is, by construction, a false positive; the procedure recovers an expected empirical FWER of 0.05.

At η^2^=0.3, an average of 252 out of 1000 parcels reach significance. These are enriched among the most extreme (highest- and lowest-ranked) parcels of the true GT network, so significance tracks genuine GT structure rather than being scattered at random (Fig. 1f).

The count, however, depends strongly on PC1 alignment: GTs strongly aligned with PC1 yield over three times as many significant parcels as weakly aligned ones (Fig. 1g), mirroring the PC1 inflation seen throughout. Parcel-level significance, therefore, partly reflects a GT’s alignment to PC1 rather than the strength of its specific signal, and should be interpreted with that dependence in mind.

The previously introduced eigencorrection of the lesions’ FC map (*corrected sLNM*) largely removes this PC1-dependence in the resulting maps: for the *corrected sLNM maps*, the significant fraction is more flat across corr(GT, PC1) (see Fig. 1g), bringing the average from 252 to 101 out of 1000 parcels. Crucially, the two curves nearly coincide at low |corr(GT, PC1)| and diverge only toward the extremes. Therefore, the excess significance at the extremes in the default maps reflects PC1-driven inflation that can be partially corrected using our *corrected sLNM* method.

## Conclusion

Our analytic and simulation framework enables formal evaluation of the claims made by both proponents and critics of LNM, and yields two main results.

First, although each sLNM map is low-dimensional (a rank-7 reconstruction captures 87% of its variance), it nonetheless recovers its own GT well above the level achievable from PC1 alone. Low effective dimensionality, therefore, does not force distinct maps to collapse onto a common pattern.

Second, the bias towards PC1 is real and has a definite source: it is inherited from the PC1-dominated lesion FC maps. A targeted flattening of the leading connectome modes for the FC construction at least halves the bias in the resulting sLNM maps and yields a recovery profile that is more uniform across corr(GT, PC1). If empirically observed symptom scores carry little PC1 bias, this modification can improve overall recovery. The correction also reduces the inflated number of significant parcels in PC1-aligned GT networks.

The simulation results depend on the structure of the GT networks used. Using the 25 networks derived from Independent Component Analysis (ICA) of fMRI data from the Human Connectome Project^10^ instead of Schaefer parcels, for example, yields substantially lower recovery for spatial r, whereas symptom r and top-100 overlap are less affected (Supplementary Fig. 3). Since ICA networks load more variance on higher connectome eigenmodes than Schaefer parcels’ connectivity profiles (see Supplementary Fig. 4), their structure is preserved less faithfully by sLNM, whose construction is dominated by the connectome’s leading eigenmodes (Supplementary Note 2). This suggests that sLNM’s spatial recovery depends on the smoothness of the underlying GT network.

Encouragingly, real sLNM maps show the same separability we predict here: projecting the set of 14 representative LNMs used by van den Heuvel et al. onto the first three principal components already separates them visibly in this low-dimensional space (Fig. 1h), consistent with our finding that low dimensionality does not necessarily lead to loss of symptom-specific information. This finding is also in line with Edelman et al.^5^.

Together, these results show that the critique raised by van den Heuvel et al. reflects genuine methodological issues that should be taken into account in future studies. However, they do not invalidate LNM as an approach. What remains untested here is whether this holds for subcortical circuit nodes, which are central to LNM’s translation into deep brain stimulation targets. Extending the present framework and the correction to these structures is a natural and necessary next step.

## Author contribution

V.L. conceived the study. T.Z. performed the simulations and analytical work under the supervision of V.L. and prepared the figures. T.Z. and V.L. wrote the paper. Both authors reviewed and approved the final manuscript.

## Code Availability

The simulation and analysis code is available on github: https://github.com/thomaszaugg/sLNM-dimensionality

## Supplementary material

**Supplementary Fig. 1:**
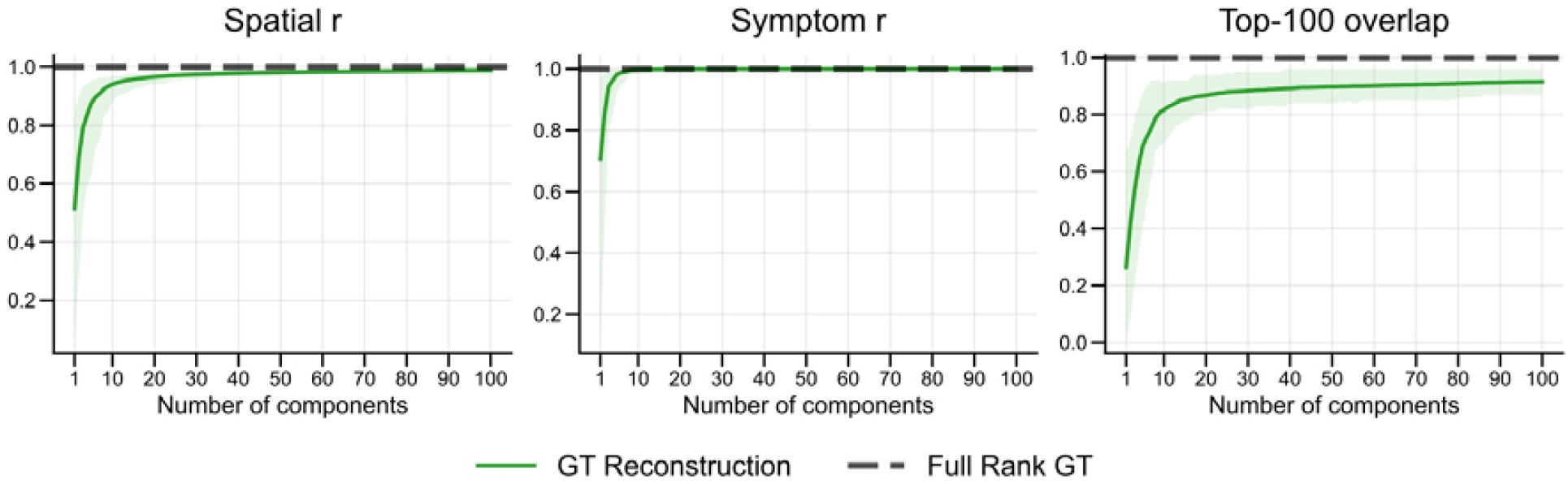
Self-recovery of lower-rank GT reconstructions. For spatial r (left) and symptom r (middle), lower-rank reconstructions need 7 and 3 principal components, respectively, to reach 90% of the full-rank GT (dashed) recovery. Top-100 overlap (right) needs 58 components to reach 90% recovery, indicating that the recovery of peak regions rests on higher components of the connectome. Error bands span the 5th–95th percentile across 300 GT networks.

**Supplementary Fig. 2:**
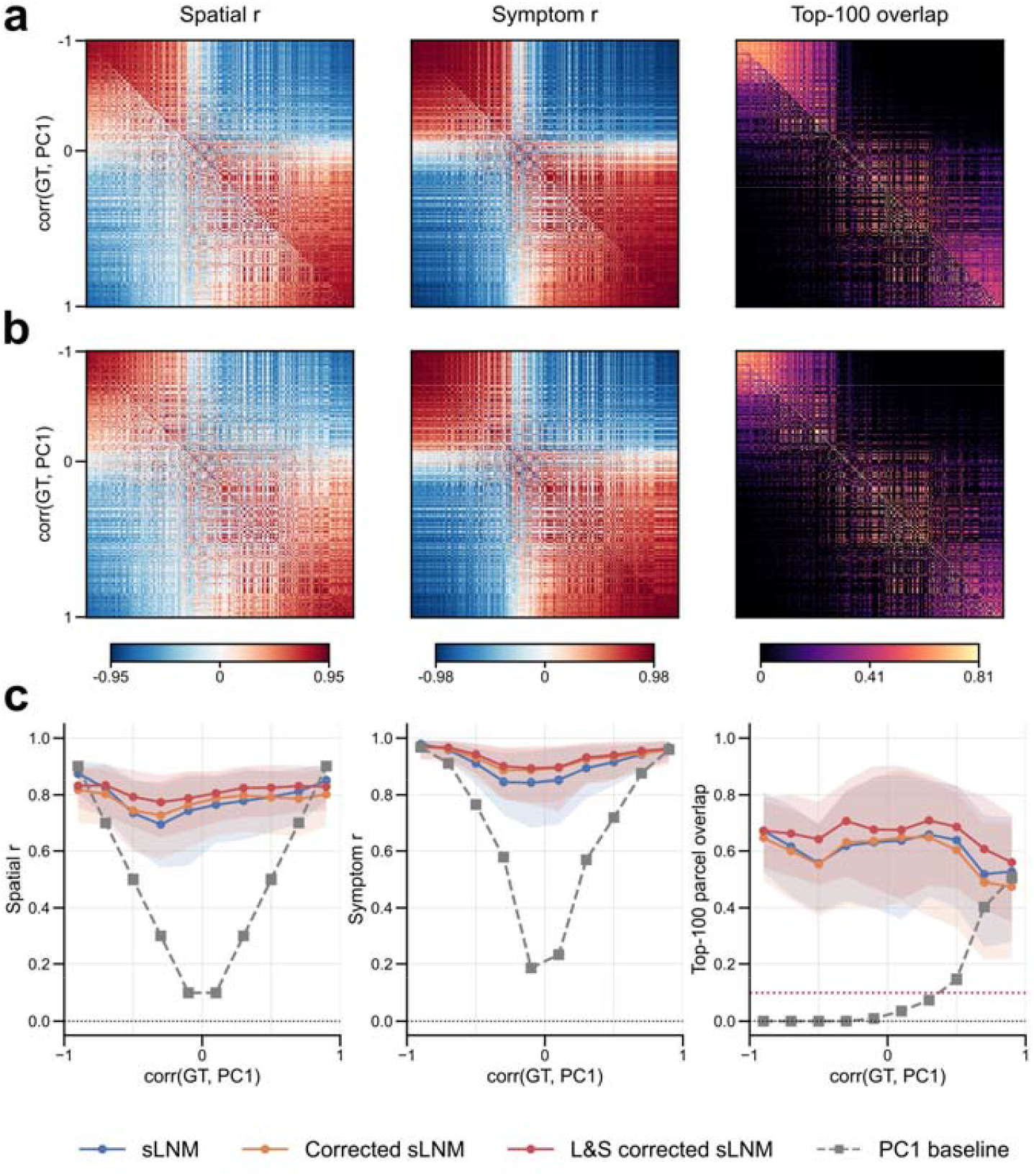
**a**,**b)** Pairwise similarity among the 300 GT networks (lower triangle) and the corresponding sLNM (upper triangle), sorted by corr(GT, PC1). First row (a) is corrected sLNM, second row (b) is L&S corrected sLNM. *Spatial r* (left): the block structure of the GT set (lower triangle) is reproduced in both corrected sLNM (a) and L&S corrected sLNM (b, upper triangle). The PC1-driven inflation found in the original sLNM (Fig. 1a) is reduced in the corrected sLNM; L&S correction shows essentially no inflation. *Symptom r* (middle) between map pairs is strongly positive or negative, tracking the sign of each pair’s PC1 alignment. *Top-100 overlap* (right) shows fine-grained spatial structure for both GT and sLNM. **c)** Recovery of GT for the original (blue), corrected (orange), and L&S corrected sLNM (red) for spatial r (left), symptom r (middle) and top-100 overlap (right). All three conditions stay well above the PC1 baseline (grey). The correction raises recovery for low-to-moderate PC1-aligned GT with respect to original sLNM (blue), with L&S corrected (red) having a higher recovery than corrected sLNM (orange). For highly PC1-aligned GT, the correction removes the recovery inflation driven by PC1 bias, leading to a flatter recovery profile across corr(GT, PC1). Error bands span the 5th–95th percentile across 300 GT networks and 10 noise realizations each; η^2^ = 0.3 throughout.

**Supplementary Fig. 3:**
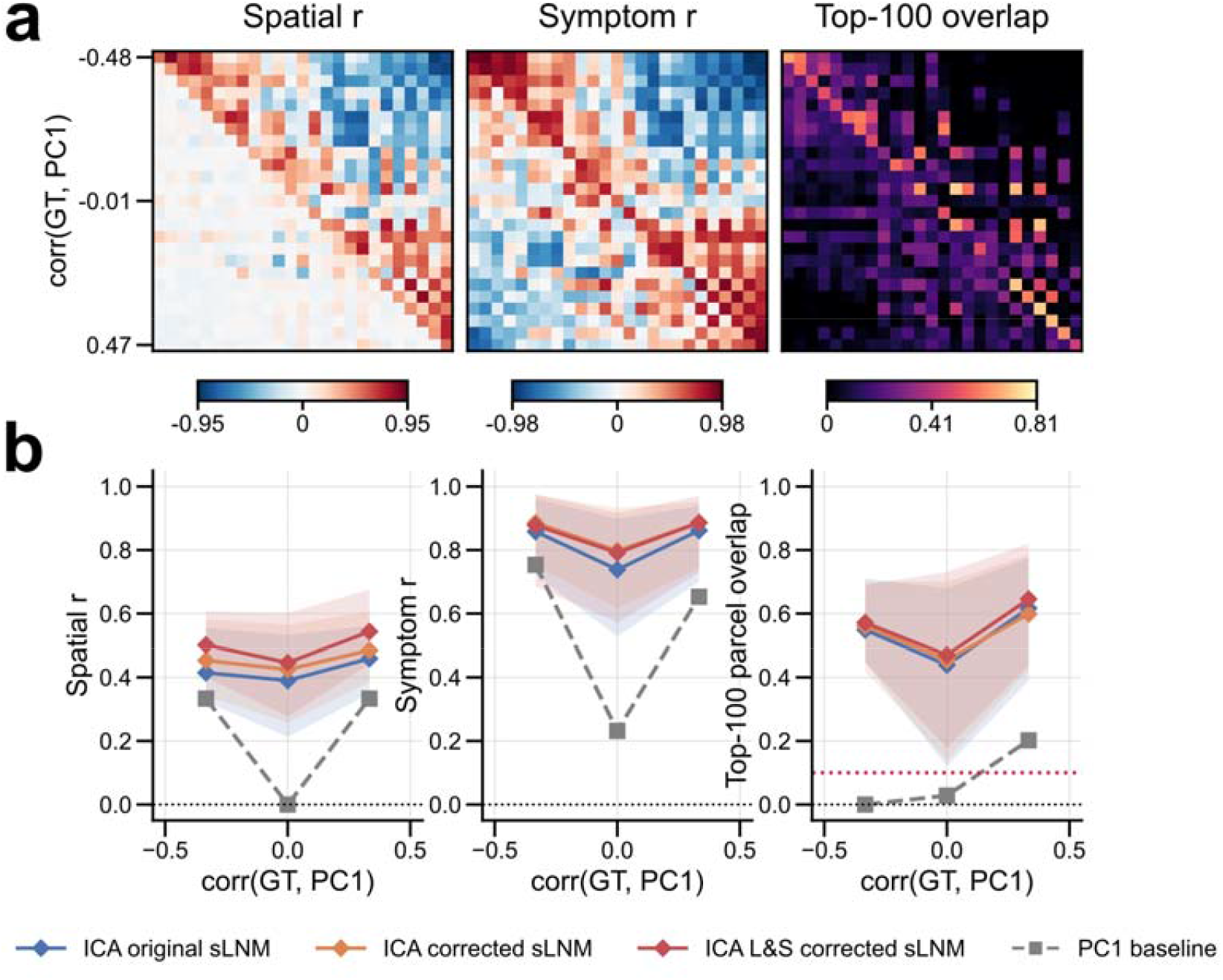
sLNM built from the 25 ICA ground-truth networks (ICA-GT) rather than the 300 parcel-derived GT networks used in Fig. 1. Conditions and triangle convention follow Fig. 1a: lower triangle, ICA-GT pairwise similarity; upper triangle, ICA original sLNM pairwise similarity; diagonal, sLNM-GT recovery for each network. **a)** Spatial r (left) shows highly inflated similarity among sLNM (upper triangle) relative to ICA-GT (lower triangle). Symptom r (middle) is polarized even among ICA-GT pairs themselves. Top-100 overlap (right) shows little overlap for ICA-GT pairs, whereas the resulting sLNM have inflated overlap. **b)** Recovery of ICA-GT for ICA original (blue), ICA corrected (orange), and ICA L&S corrected (red) sLNM, against the PC1 baseline (gray), as in Fig. 1c. Spatial r recovery is roughly halved relative to parcel-GT (Fig. 1c), while symptom r remains high and top-100 overlap is only slightly reduced, indicating that peak regions remain largely recovered despite the drop in overall spatial correlation. The correction raises recovery throughout, with ICA L&S corrected sLNM having higher recovery than ICA corrected sLNM. Despite the drop in recovery, sLNM all have higher recovery than the PC1 baseline. Error bands span the 5th–95th percentile across 25 ICA-GT networks and 10 noise realizations each; η^2^ = 0.3 throughout.

**Supplementary Fig. 4:**
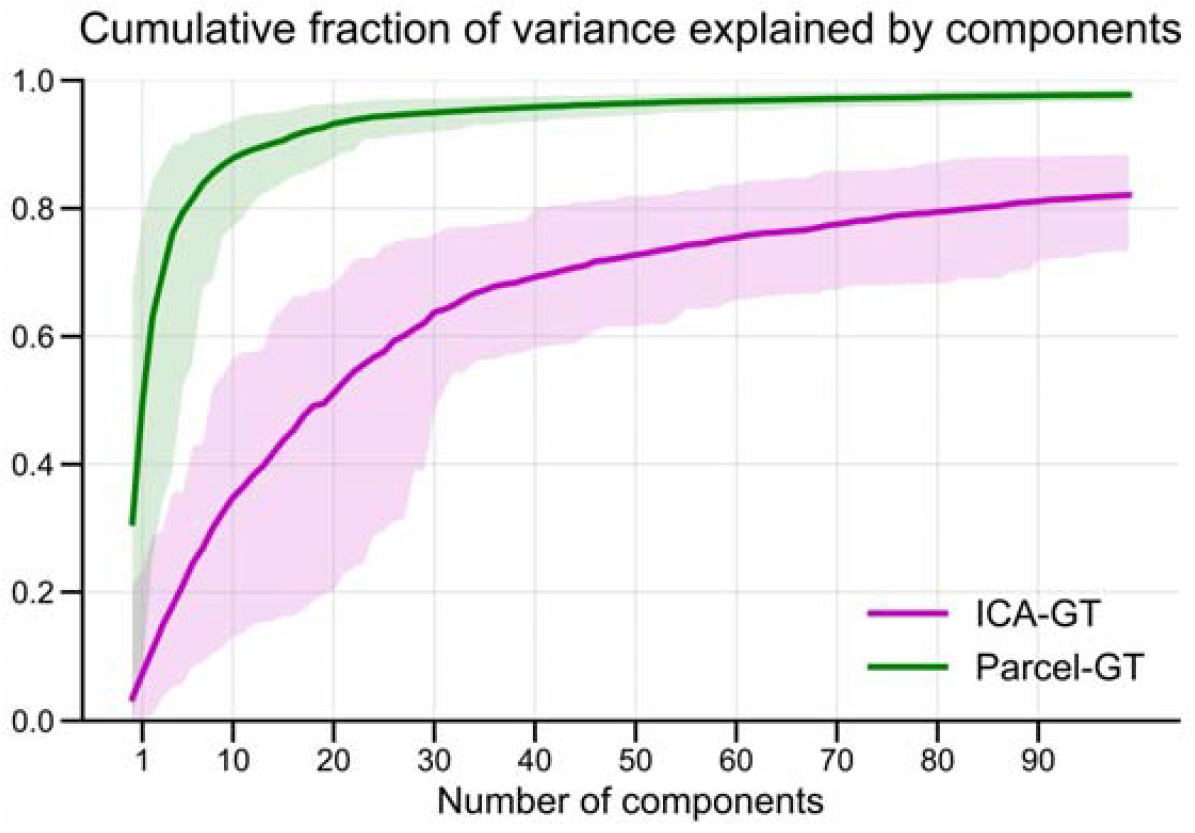
Cumulative fraction of variance explained as a function of the number of components, for GT networks derived from Schaefer parcels (parcel-GT, green) and from ICA components (ICA-GT, magenta). Parcel-GT reaches a cumulative variance of 0.8 within the first 6 components, whereas ICA-GT only reaches with 86 components, indicating that ICA-GT variance loads onto higher-order connectome eigenmodes than parcel-GT. Error bands span the 5th–95th percentile across 300 parcel-GT networks and 25 ICA networks, respectively.

**Supplementary Fig. 5:**
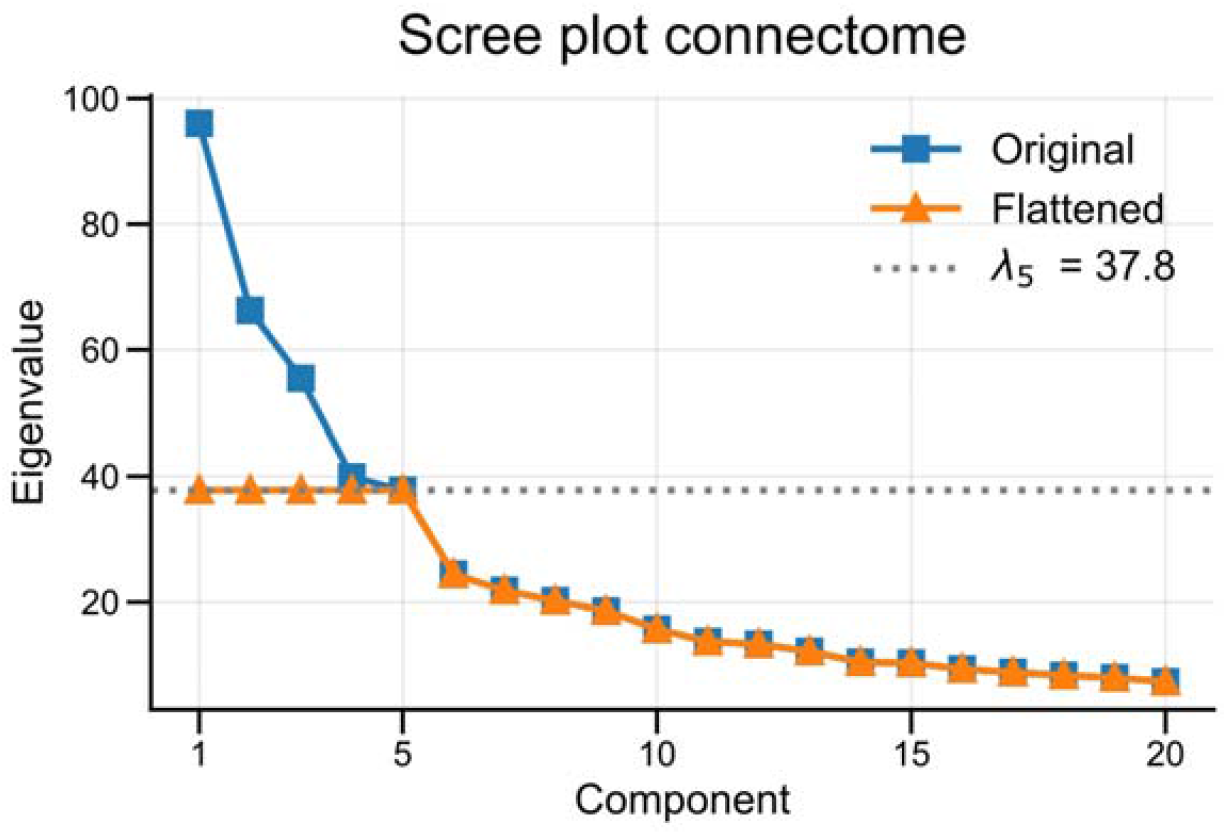
Scree plot of the GSP1000 connectome before (blue) and after (orange) flattening the leading four eigenvalues to the level of the fifth (λ_5_, dotted), the spectral elbow. Components 6 and beyond are unchanged.

**Supplementary Table 1.** Real LNM maps referenced in Fig. 1h.

| Fig. 1h label | Disease network | Filename in Repository* | Reference |
| --- | --- | --- | --- |
| 1 | Epilepsy | JI_EPILEPSY | Ji, G.-J. *et al.* A generalized epilepsy network derived from brain abnormalities and deep brain stimulation. *Nat Commun* **16**, 2783 (2025). |
| 2 | Tremor (DBS) | JOUTSA_TREMORDBS | Joutsa, J. *et al.* Identifying therapeutic targets from spontaneous beneficial brain lesions. *Annals of Neurology* **84**, 153–157 (2018). |
| 3 | OCD (DBS) | LI_OCDDBS | Li, N. *et al.* A unified connectomic target for deep brain stimulation in obsessive-compulsive disorder. *Nat Commun* **11**, 3364 (2020). |
| 4 | Psychosis | PINES_PSYCHOSIS | Pines, A. R. *et al.* Mapping Lesions That Cause Psychosis to a Human Brain Circuit and Proposed Stimulation Target. *JAMA Psychiatry* **82**, 368–378 (2025). |
| 5 | Anxiety | SIDDIQI_ANXIETYi | Siddiqi, S. H. *et al.* Distinct Symptom-Specific Treatment Targets for Circuit-Based Neuromodulation. *Am J Psychiatry* **177**, 435–446 (2020). |
| 6 | Depression | SIDDIQI_DEPRESSION | Siddiqi, S. H. *et al.* Brain stimulation and brain lesions converge on common causal circuits in neuropsychiatric disease. *Nat Hum Behav* **5**, 1707–1716 (2021). |
| 7 | Depression (MS) | SIDDIQI_MSDEPRESSION | Siddiqi, S. H. *et al.* Lesion network localization of depression in multiple sclerosis. *Nature Mental Health* **1**, 36–44 (2023). |
| 8 | Political behaviour | SIDDIQI_POLITICAL | Siddiqi, S. H., Balters, S., Zamboni, G., Cohen-Zimmerman, S. & Grafman, J. H. Effects of focal brain damage on political behaviour across different political ideologies. *Brain* **148**, 3280–3289 (2025). |
| 9 | PTSD | SIDDIQI_PTSD | Siddiqi, S. H. *et al.* A potential target for noninvasive neuromodulation of PTSD symptoms derived from focal brain lesions in veterans. *Nat Neurosci* **27**, 2231–2239 (2024). |
| 10 | Criminal behaviour | DARBY_CRIMINALITY | Darby, R. R., Horn, A., Cushman, F. & Fox, M. D. Lesion network localization of criminal behavior. *Proceedings of the National Academy of Sciences* **115**, 601–606 (2018). |
| 11 | Tics | GANOS_TICS | Ganos, C. *et al.* A neural network for tics: insights from causal brain lesions and deep brain stimulation. *Brain* **145**, 4385–4397 (2022). |
| 12 | Addiction | JOUTSA_ADDICTIONCONTRASTAB | Joutsa, J. *et al.* Brain lesions disrupting addiction map to a common human brain circuit. *Nat Med* **28**, 1249–1255 (2022). |
| 13 | Aphantasia | KUTSCHE_APHANTASIA | Kutsche, J. *et al.* Lesions causing aphantasia are connected to the fusiform imagery node. *Cortex* **197**, 1–12 (2026). |
| 14 | Aphasia | ZARIFKAR_APHASIA | Zarifkar, P. *et al.* Lesion network mapping of eye-opening apraxia. *Brain Commun* **5**, fcad288 (2023). |

\* https://github.com/dutchconnectomelab/lesionnetworkmapping

## Supplementary Note 1: Simulation

Following Treeratana et al., we used the GSP1000 group-average matrix as the functional connectome^11^. Its leading eigenvector (PC1) closely tracks the connectome degree map (r = 0.86) and dominates a strongly top-heavy spectrum (Supplementary Fig. 5).

Each of the 300 ground-truth (GT) networks is the whole-brain functional connectivity map of a single cortical parcel. As this atlas was designed to maximise between-parcel distinctiveness in connectivity profiles^7^, it is well suited to this purpose.

Each lesion is a 4-mm-radius sphere placed at a randomly sampled brain voxel and represented by its voxel-wise functional connectivity (FC) map, computed as seed-based temporal average Pearson correlation (see Treeratana et al.^6^). For our simulation, the lesion FC maps are projected to parcel space by averaging within each Schaefer-1000 parcel.

A lesion’s symptom signal is determined by how well its FC map matches the GT (the Pearson correlation r_GT between them). Observed symptom scores combine this signal with Gaussian noise at a signal fraction η^2^, the proportion of symptom variance attributable to the GT:

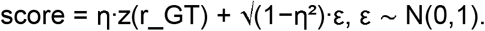

We set η^2^=0.3 following the *weak effect size* of Treeratana et al.^6^.

Each sLNM map is built from 100 lesions bootstrapped with replacement from the 500-lesion pool, as in Treeratana et al., with the symptom scores computed as explained above. We follow the simplified sLNM method as proposed by van den Heuvel et al.: the sLNM map is the parcel-wise correlation, across lesions, between each lesion’s FC value at this parcel and the respective symptom scores ^1^. We repeat this for all 300 GT networks across 10 noise realisations each.

We compute sLNM maps in parcel rather than voxel space. The two approaches are near-identical (mean r = 0.99 after projection), so this introduces no meaningful methodological divergence from Treeratana et al.

## Supplementary Note 2: PC1 alignment

The source of the PC1 bias lies in the lesions’ functional connectivity (FC) maps. A lesion’s FC map is its seed passed through the connectome:

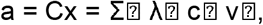

where C is the functional connectome, x the lesion seed, λ- and v- the eigenvalues and orthonormal eigenvectors of C, with v_1_ corresponding to PC1. The loading c- = v-□ x is the projection of the lesion seed onto the m^th^ eigenmode. Each loading c- is scaled by its eigenvalue λ-, so the strongly top-heavy spectrum amplifies the leading modes. A seed only modestly aligned with PC1, therefore, yields an FC map dominated by it.

For example, consider a lesion that is loading more on the fifth eigenmode than the first. With loadings c_1_ = 0.6 and c_1_ = 0.8 and the empirical λ_1_ = 96, λ_5_ = 38, the FC map carries 0.6 · 96 = 57.6 on PC1 against 0.8 · 38 = 30.4 on v_5_. The lesion FC map is PC1-dominated even though the seed was more aligned with v_5_.

## Supplementary Note 3: Corrected sLNM maps

We correct the PC1 bias by flattening the top eigenvalues of the connectome spectrum before building the lesion FC map. The connectome is low-dimensional; therefore, most eigenmodes carry little signal. In the original sLNM, the eigenvalue weighting already suppresses noise-dominated tail modes on its own. Our proposed correction touches only the leading modes, where most of the inflation occurs, and leaves the remainder unchanged. The optimal normalisation is connectome-specific and reflects a trade-off between attenuating PC1 alignment and preserving SNR; we flatten modes 1–4 to the level of mode 5, the elbow of the GSP1000 eigenspectrum (Supplementary Fig. 5).

Lesions’ functional connectivity is computed in voxel space, and only projected to parcel space afterwards; the voxel-wise connectome itself is never explicitly formed, so flattening its spectrum directly is not possible.

The correction can nonetheless be applied after projection: the lesion’s parcel-space FC map is projected onto the connectome’s parcel-level eigenvectors, the loadings of modes 1– 4 are rescaled by w- = λ_5_/λ-, bringing their effective eigenvalues down to that of mode 5, and the corrected map is reconstructed into parcel space from these rescaled coefficients.

The same principle extends to voxel space: computing the first four voxel-wise eigenvectors and eigenvalues, projecting the lesions onto them, and rescaling before reconstruction achieves the correction at the voxel level.

## Supplementary Note 4: Comparison with Treeratana et al.’s null and correction

Treeratana et al.’s null addresses the comparison of two sLNM maps^6^. They evaluate the spatial r between two maps against the distribution of spatial r obtained after permuting symptom labels, testing whether two datasets converge more than chance predicts. This is a global convergence statistic for a map pair.

Our null addresses a different question, at the level of a single map. For each map, we permute symptom labels and compare each parcel’s observed value against the null distribution of the maximum absolute parcel value across the map, which controls the family-wise error rate over all 1000 parcels. This yields a per-parcel significance map rather than a single global statistic.

To prevent the over-similarity of two maps, Treeratana et al. propose to regress PC1 out of the sLNM maps before testing convergence. This successfully prevents the over-convergence of sLNM map pairs. We propose, on the other hand, to correct the PC1 bias during the lesions’ FC construction. This not only reduces the PC1 bias for subsequent test statistics, but also corrects the map itself.

